# Molecular phenotypes that are causal to complex traits can have low heritability and are expected to have small influence

**DOI:** 10.1101/009506

**Authors:** Leopold Parts

## Abstract

Work on genetic makeup of complex traits has led to some unexpected findings. Molecular trait heritability estimates have consistently been lower than those of common diseases, even though it is intuitively expected that the genotype signal weakens as it becomes more dissociated from DNA. Further, results from very large studies have not been sufficient to explain most of the heritable signal, and suggest hundreds if not thousands of responsible alleles. Here, I demonstrate how trait heritability depends crucially on the definition of the phenotype, and is influenced by the variability of the assay, measurement strategy, and the quantification approach used. For a phenotype downstream of many molecular traits, it is possible that its heritability is larger than for any of its upstream determinants. I also rearticulate via models and data that if a phenotype has many dependencies, a large number of small effect alleles are expected. However, even if these alleles do drive highly heritable causal intermediates that can be modulated, it does not imply that large changes in phenotype can be obtained.

## Introduction

High heritability of human features has now been observed over centuries (e.g. [1,2]). One natural research program for understanding and eventually modulating such traits, including common disease susceptibility, focuses on finding and analyzing their causal intermediates. These phenotypes lie in the chain of biochemical reactions that produce the complex trait, and can effect it when perturbed. The premise is that by creating a large compendium of the intermediate measurements, and mapping their genetic basis, we can identify ones that share associated alleles with a disease, and are thus likely candidates to be the causal mediators. Perhaps they can also be directly modulated, yielding a promising drug target.

So far, studies in this direction have focused mainly on gene expression, and to a lesser extent, protein abundance and metabolite levels as candidate phenotypes, as these traits can be measured comprehensively in high throughput manner. A further argument for this choice is that stronger genetic signal is expected to be carried by immediate downstream products of the DNA sequence. Both segregation of molecular phenotypes based on the complex disease status, as well as co-localised genetic control of the two have been sought for and observed [3]. However, studies put the median mRNA heritability estimates in human blood to only ∼10–20% [4–6], leaving a large gap with disease heritabilities (e.g. ∼40% for type two diabetes [7]).

In the following, I will first show that the observed low heritability of molecular traits can theoretically and practically be increased by fixing, subtracting, or averaging over confounders. However, mediators of the genetic signal of highly heritable complex traits do not necessarily have to be highly heritable themselves; heritability can increase as the considered trait is more downstream from DNA. I also present a generative model for phenotypes that predicts many small effect alleles without requiring influential mediators, which is consistent with a range of observations of molecular and organism traits.

## Definition of the phenotype is central to heritability estimation

As a motivating example, consider the discrepancy between high disease risk, and low RNA abundance heritabilities. What should one expect for protein levels that mediate between the two? One recent study measured protein abundance in large populations of genetically diverse single cells by fluorescence [8]. The heritability of protein level can be estimated in two ways from these data. First, one can use comparisons of population average values between individuals of different degrees of relatedness to calculate the genotype effect, and repeat measurements to establish the extent of random environmental variability. If the replicates are highly concordant, the non-genetic contributions explain only a moderate part of the total variation (Fig. 1A). As a result, the distribution of heritability estimates spans the entire range from 0 to 1, with a median of 50% (Fig. 1B). Second, the phenotype can equally well be chosen to be protein abundance in a single cell. In this case, the cell to cell variance component dominates both genotype signal and repeat variability (Fig. 1C), leading to lower heritability estimates (median 3%, Fig. 1D).

**Figure 1.**
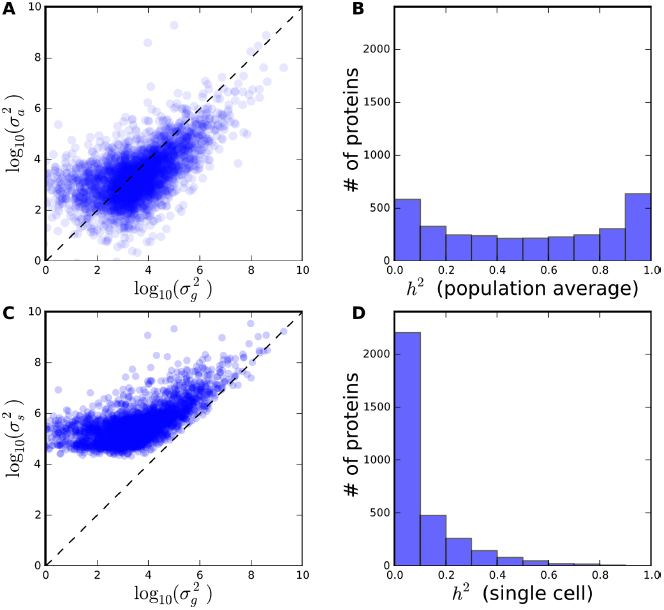
Heritability estimate depends on phenotype definition via the total phenotypic variance. Genetic (*σ*^2^_*g*_) assay (*σ*^2^_*a*_) and single cell (*σ*^2^_*s*_) variance of population average abundance in a yeast cross (data from [8]). A. Genetic and assay variance of protein level. Every dot corresponds to one GFP fusion strain for which fluorescence was measured in ∼50,000 clonal and segregating cells to derive the variance estimates. B. Distribution of heritability estimates *σ*^2^_*g*_/(*σ*^2^_*g*_ + *σ*^2^_*a*_ + *C*^−1^*σ*^2^_*s*_) of population average protein abundance. C. Genetic and non-genetic variance of single cell protein abundance; see (A). D. Distribution of heritability estimates *σ*^2^_*g*_/(*σ*^2^_*g*_ + *σ*^2^_*a*_ + *σ*^2^_*s*_) of single cell protein abundance.

A simple redefinition of the studied phenotype from single cell measure to a population average has a drastic effect on the estimated heritability. We can examine how this emerges under the general linear model framework that decomposes a trait into all the different contributing sources (e.g. [9,10], Box 1). A heritability estimate compares the magnitude of the genotype effect to all the sources of variation combined, and the latter can be influenced by choices in experimental design and analysis. First, if a confounding effect (such as assay batch, smoking status, or gender) is fixed, it cannot introduce variability in the trait. Second, defining a phenotype as an average over any of the non-genetic sources of signal reduces the corresponding variance component, leaving the others constant. For example, if variance due to the assay replicate is retained, the total variance is larger, and heritability thus smaller compared to the phenotype averaged across replicates. If we knew the full generative model of the trait, we can integrate over all the sources of confounders that comprise the non-genetic contribution, and make the genotype signal arbitrarily large relative to the total variation. Third, unobserved sources of variability can be modeled using principal components or factor analysis [10,11], which is approximately equivalent to normalising for their effect. The heritability estimates for high-dimensional molecular traits (e.g. [12,13]) can increase considerably if the total trait variance is reduced by correctly subtracting effects of unobserved confounders [14], but also decrease if the components carry genetic signal [15]. Finally, data normalisation and transformation will influence the heritability estimates. A principled optimisation of the transformation (such as quantile normalisation, taking logarithms, etc.) applied to raw data can increase accuracy of parameter inference and mapping power [16]. Departures from the underlying assumptions of linear models, and conversely, data transformations to satisfy them, will influence the heritability estimates as well.

### Box 1. Molecular trait heritability under different definitions

*Average trait from nested observations.* Let trait *y* be observed in individual *i*, replicate *j*, in *c* cells, with average value *μ*, and random effects only due to genotype *g*, assay *a*, and single cell 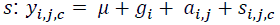, or 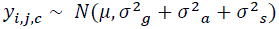. The standard definition of heritability is the fraction of total variance that is genetic: 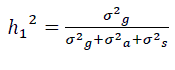. Now, consider the average of *y* across *C* cells and *J* assay replicates: 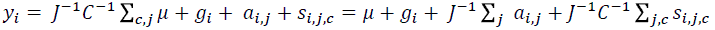, with heritability estimate 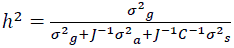. In this reformulation of the trait, all non-genetic sources of variation are averaged over, so their contributions to the total variance are decreased, increasing the relative strength of genotype signal. If number of replicates *J* is large, the heritability will increase to 1. Statistically, we are replacing the variance of the confounder with the sampling variance of its mean, which can be reduced by making additional observations. Traits can also be redefined by fixing covariates, subtracting effects of unobserved confounders, and transforming the measurements as discussed in Text and Supporting Text.

*A combination of multiple different traits.* Let *x*_1_ and *x*_2_ be traits with equal variances and heritabilities. If their genetic contributions are positively correlated, but environmental influences independent, the genetic signal is aggregated from the two traits, and the heritability of *y*=*x*_1_+*x*_2_ is larger than that of *x*_1_ or *x*_2_. In general, the heritability of a weighted sum of traits 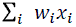 is the weighted harmonic mean *H* of their heritabilities and genetic covariances, scaled by the fraction of total variation that is not due to environmental covariance 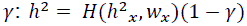. The resulting value can be both above and below the individual trait heritabilities, depending on the correlation structure between the genetic and environmental influences. The explicit derivations of results in Box 1 and 2 are given in the Supporting Text.

These steps are not just mathematical reformulations of the data, but define what the phenotype means, which is then reflected in the heritability estimate. A disease trait can be interpreted as a label of having a disease, a personal risk probability, or a normal liability score; for molecular traits, one can consider instantaneous rates, cumulative levels, time- and space averages, and it is not immediately obvious which choice is most appropriate. From heritability estimation perspective, the redefinitions of phenotype can equivalently be viewed as obtaining a new population in which some chosen influences do not introduce trait variability. As heritability is a population statistic, it is necessarily altered under such changes. Thus, to interpret any estimate in the literature, it is essential to understand what constitutes the population, or equivalently, which sources of phenotypic variance are included and excluded from the heritability calculations. If the goal of the study is to understand the effect of genotype on a trait using mapping, it is useful to optimize the trait heritability by redefining the phenotype to reflect the true parameter of interest without the confounding influences.

## Mechanism of the phenotype can alter the strength of genetic signal

While changing definitions can alter our view of trait heritability, there is still a reasonable intuition that all other factors being equal, heritability of a downstream trait cannot be larger than an upstream one (e.g. [17]). Additional layers of regulation should only increase variability, but not make genetic signal appear from nowhere. However, this does not hold in general for traits that have multiple upstream factors, as it is possible to combine their genetic influences. For example, the human height phenotype is a sum of many individual bone lengths, and some additional factors. Height heritability has been estimated to be around 80% [2,18], and even over 90% in some populations [18,19], while estimates for individual bone length are slightly smaller (57–73%, [20]). Theory and data support shared control of different bone lengths via hormone and growth factor abundance during development [21,22], some of which is genetic. All environmental influences do not have to be completely correlated, as different bones obtain their length at different times [23].

The heritability of a phenotype that is a combination of multiple traits can extend beyond the heritabilities of its constituents (Box 1). For example, if there is no correlation between the random genetic or environmental effects influencing a set of traits, the heritability of their sum lies between the heritabilities of the summands. However, if the traits share causal genetic factors, but environmental influences are independent, the sum of the traits can have a relatively stronger genetic contribution, as the uncorrelated environmental signal is averaged away. The intuitive reason is the same as for removing replicate variance: the genotype signal is integrated over the noisy measurements. Notably, heritability can also decrease if genotype contributions are negatively correlated due to antagonistic pleiotropy.

The idea of combining noisy phenotypes to precisely reflect the underlying genetic cause can also be used to derive a more accurate upstream phenotype from many downstream ones. The effect of a common regulator that induces correlation between traits can be recaptured by averaging out the additional independent variance (e.g. [24–27]). The value of such an approach was demonstrated on a large-scale genetics of gene expression dataset from three tissues in British twins. The authors inferred pathway phenotypes from the expression data by combining signal from multiple genes in a pathway to five quantitative pathway activity traits. The resulting variables were highly heritable, associated with age, and could be easily interpreted, as they were derived from biological prior knowledge [28]. The increased heritability is consistent with shared genetic basis of pathway expression regulators or pathway members themselves, and could allow more powerful genetic mapping [24,25].

## Simple biochemical models predict polygenic traits with few influential intermediates

Regardless of the exact definition of heritability, it is clear that most traits have a genetic component, and there are alleles to be mapped for the causes. So far, the combined contribution of identified associated variants has been substantially smaller than the total amount of heritable variation. Reasons proffered to account for this discrepancy range from inaccurate heritability estimates due to epistasis [29], synthetic associations [30], and many rare alleles of large effect [31] to a large number of common variants with small effects [32,33]. We can ask which of these are concordant with different models of the genotype-phenotype relationship and its evolution [34] while more data are gathered from even larger cohorts for explicit tests.

A natural model based on our physical understanding of the world considers any phenotype as a product of chains of biochemical reactions. Under some simplifying assumptions, such as free diffusion and law of mass action, this description can be made explicit, and used to derive analytical results (Box 2). A single reaction takes place with a characteristic rate, which can be interpreted as an inverse of a timescale. The timescale of one reaction chain is then the sum of the timescales of individual reactions, coupling their influences [35,36]. This interdependence between many steps implies that the majority of them can only have limited contribution to the downstream product, suggesting the existence of many small effects. Further, the influence of the speed of any single step, and therefore its share of the total variation, is inversely proportional to the activity of the responsible proteins. Thus, the presence of two copies of a gene in humans explains the widespread recessivity of individual alleles and additive effects [36]. This model also predicts widespread functional epistasis, both within a gene as recessivity, as well as between genes [37,38], which has been observed in screens and crosses of model organisms [39–43].

### Box 2. Enzyme-inspired model of a complex trait (after Kacser and Burns).

Consider a pathway for a molecule *X*_*n*+1_ that starts with a source molecule *X*_0_, and is produced via intermediates *S*_1_ to *S_n_* with the action of proteins *E*_0_ to *E_n_*.

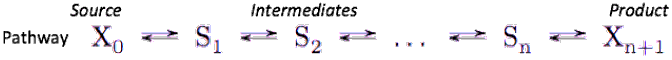

Each of the *n* + 1 steps has a characteristic timescale *τ*. that is inversely proportional to the total activity of the protein [*E*_*i*_] from the two copies in the genome, 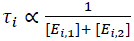. The complex trait *F* is the change in *X*_*n*+1_ over time, which is inversely proportional to the total timescale of reaction chain, 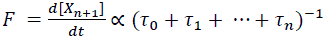. The sensitivity of *F* to any individual protein *E*_*i*_ is limited due to coupling with the other steps, 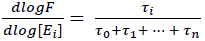. Even if [*E_i_*] itself has heritability of 1, the influence of its modulation on the phenotype is restricted to how sensitive *F* is to it. This treatment of a single chain can be extended to include more general topologies, such as branching [47]. Of course, insights from such an abstraction are contingent on the model assumptions reflecting reality, and the applicability has been discussed in different venues (e.g. [36,37,48]).

Under this view of complex traits, the effect of a causal allele can be separated into its contribution to a protein activity, and the sensitivity of the phenotype to that protein. The abundance and activity of the protein itself are a result of several processing steps from transcription, posttranscriptional modifications, nuclear export, translation, and posttranslational processing. Each of the required steps can have several associated alleles segregating in the population, explaining some of the existing variation, but limited in total contribution to the trait heritability. This suggests that there can be many *trans*-acting alleles, but the relative importance of any one is restricted. Consistent with this model, the genetic control of molecular traits can be substantial. *cis*-acting variants with few protein intermediates and direct effect on transcription have been shown to explain a majority of variation of some transcript levels in model organism crosses, as well as outbred populations and human cohorts [44,12]. *trans* effects that operate on protein levels indirectly are found to be more frequent and estimated to be smaller [45,8]. The risk of incidence, and onset characterisation of disease phenotypes have many upstream contributors, each influenced by activities of several proteins, that further vary due to effects of multiple alleles in the population. As the sensitivity of phenotype to intermediate protein activity has to be low for most proteins, modulating even the highly heritable intermediates that truly causally influence the organism phenotype does not have to lead to a large effect. Abundant small effect mutations are therefore expected for all traits that are products of many genes. Even if major genes with larger effects exist, such as rate-limiting steps in pathways (e.g. [46]), many modifiers are predicted.

## Discussion

I put forth a statistical and a biological reason why heritability estimates for organism traits can be higher than for their causal molecular phenotypes. From purely numerical considerations, the molecular trait could be defined inappropriately. The genotype signal is present in measurements confounded by stochasticity, timing, location, and assays, but in order for it to have a large relative impact, the phenotype has to be redefined as an integral over the other sources of variation. Any measurement can in principle be used for heritability estimation, thus leaving a choice as to how many different axes of variability to average out, and thereby, how large of a heritability to obtain. A similarly inappropriate definition is positing a standard linear model on a non-Gaussian distribution, which can also reduce the heritability estimate. Biologically, the phenotype itself can be a product of several lower traits which are more noisy, but share some genetic basis. The combination of multiple correlated genetic signals strengthens it, and results in a trait that truly is more heritable than any of its upstream causes. I demonstrated analytically (Supporting Text) that the resulting heritability of a linear combination of traits is the weighted harmonic mean of constituent heritabilities and genetic covariances.

These results caution against taking heritability estimates of molecular traits at face value, especially for comparisons across multiple studies with different sampling designs and phenotype definitions. They also suggest a tension between integrating signals from genotype and environment over a long time period. Disease risk and many other life-history traits are highly heritable [49], but the estimates can both increase and decrease over time. For example, the heritability of human body-mass index increases from youth to early adulthood, and then decreases again [50], suggesting a correlated genotype influence in one period, but sustained environmental effect in another. Any attempt at explaining these phenomena via heritability and genetic basis of molecular phenotypes must be careful in relevant definitions and their interpretation.

Is heritability a useful concept for understanding the molecular basis of complex traits at all? Knowledge of narrow sense heritability in a population constrains the set of possible allele effect size distributions, and puts an upper limit on the ability to map and predict phenotype using independent locus information. These can be informative for research on short-term evolution and breeding, but say little about biology. Quantitative models of phenotypic variation should aim to infer interpretable effect sizes of all sources of variability in the experiment, including additive locus effects, dominance, interactions, and confounders. It is the effect sizes that reveal genetic architecture, give relative strengths of different signals, form predictive models, and guide followups. While easier said than done, this approach has more chance of furthering our understanding of what determines the phenotype. Heritability estimates and p-values can remain post-hoc checks of how much additive variation is unaccounted for, and how extremely the observations violate assumptions of the postulated null model.

The generative model of an organism phenotype via long chains of reactions suggests many alleles of small effect. This is a consequence of many constituent pathway members, recessivity of null and dose-modulating alleles, epistatic coupling of the individual steps, and the limited observed effects of mutations on protein abundance. The model is consistent with current evidence of lack of substantial low frequency variant signal increase in population isolate studies [51], negative selection due to greatly reduced fecundity of strong disease-predisposing allele carriers [52], overlap of complex trait loci with related Mendelian disorder genes [53], and lack of pedigree signal consistent with near-Mendelian segregation patterns [34]. On the positive side, the model predicts convergence of GWAS hits on the pathways involved, as has been the case for Crohn's disease [54], inflammatory bowel disease [55], and schizophrenia [56]. Also, longer genes are a larger mutational target, and therefore predicted to harbour more hits under the “polygenic dust” model, as was observed for schizophrenia [56]. While all consistent explanations (e.g. [29,33,57]) for the current inability to explain narrow sense heritability apply at some frequency, a large number of small effect alleles that truly contribute to the underlying biology ought to be the default expectation.

The explicit models of genotype effect imply experimental limitations to take into account. If the aim of the study is genetic mapping, the traits ought to be defined in a way that maximize narrow sense heritability, both via controlling and averaging confounders, as well as applying a suitable transformation to the data. The fewer potential upstream causes a trait has, the larger the potential for any one allele to explain a substantial portion of its variation in an outbred population. Therefore, studies that minimize the number of biochemical reactions between the applied perturbation and the phenotypic readout have a better chance uncovering large effects, which are also easier to ascertain, interpret, and validate. If a trait is downstream from DNA, then coupling of the many different influences buffer the effect of any one allelic perturbation, rendering genetic modulation difficult even when the intermediate causes are highly heritable. A potentially fruitful strategy is to find ways to combinatorially target multiple molecules in the same cell [58]. It is only a hope that a single experiment will identify a few variables that explain much of the behaviour in a large and complicated molecular system that produces a complex trait. Hope is not a strategy - the advances ahead require quantitative characterisation of individual reactions and system components in controlled settings.

## Acknowledgements

I thank Andrew Brown, Ishaan Gupta, Ben Lehner, Jared Simpson, and Oliver Stegle for feedback and discussions. LP is a Marie Curie International Outgoing Fellow, and a Canadian Institute for Advanced Research Global Scholar.

